# Reducing competition between *msd* and genomic DNA significantly improved the editing efficiency of the retron editing system

**DOI:** 10.1101/2024.06.04.597346

**Authors:** Yuyang Ni, Yifei Wang, Xinyu Shi, Qingmin Ruan, Tian Na, Jin He, Xun Wang

## Abstract

A retron is a distinct system encoding reverse transcriptase and a unique single-stranded DNA/RNA hybrid called multicopy single-stranded DNA (msDNA). The ability of msDNA to serve as a homologous recombination donor for gene editing has attracted great interest. However, the mechanism by which msDNA expression affects editing efficiency remains unclear. In this study, we show that an increase in *msd* number increased msDNA yield but was not necessarily accompanied by an increase in editing efficiency. Mechanistic studies indicate that *msd* and genomic regions competed for msDNA during recombination. As the number of *msd* increased, the amount of msDNA allocated to the genomic targets decreased, resulting in a decrease in editing efficiency. Finally, we reduced *msd* editing by expressing msDNA corresponding to the plasmid replication leading strand sequence, thus constructing a retron-based gene editing system that achieved 100% editing efficiency in the shortest time reported to date. The above results reveal a completely different features between retron-based gene editing system and oligonucleotide-mediated gene editing system and will provide theoretical guidance for the design and application of the retron system.

## Introduction

Retron is a unique reverse transcription system present in various microbial genomes (1-3). For example, the retron Ec86 system was originally discovered in *Escherichia coli* strain BL21 and consists of non-coding RNA (ncRNA) and reverse transcriptase (RT). The ncRNA contains two regions: msr and msd, where msd can directly synthesize multiple copies of single-stranded DNA (msDNA) through RT (4). The *msd* in the retron operon is the template for msd synthesis and is also the region where exogenous sequences are inserted during gene editing (5).

Since the discovery that the msDNA produced by the retron system can serve as a recombination template, the retron system has attracted widespread attention in the field of gene editing (5-8). The “annealing-integration” model is generally considered to be the msDNA-mediated gene editing mechanism (9-11). During the editing process, msDNA is integrated into the lagging strand of the DNA replicon in the form of Okazaki fragments, forming wild-type genomes and heterologous double-stranded genomes. The bacterial cell with the heteroduplex genome then undergoes another DNA replication and cell division, producing a wild-type genome and a mutant genome.

Researchers have made consistent efforts to improve the efficiency of retron-mediated gene editing. The gene editing efficiency of the first generation retron system in *E. coli* DH5α strain is only about 6*10^-4^ (5). In the second generation, Simon *et al*. improved the editing efficiency by 100-fold to 6*10^-2^ by introducing mutations in the 3′-5′exonuclease ExoX and mismatch repair protein MutS into the *E. coli* Top 10 strains (12). In the third generation, Schubert *et al*. knocked out *mutS*, 5′-3′ nucleic acid exonuclease gene *recJ*, and 3′-5′ nucleic acid exonuclease gene *xonA* in the EcNR1 strain, and changed the single-stranded annealing protein Beta recombinase to CspRecT, which significantly improved the editing efficiency to nearly 100% (13). Additional studies have shown that knocking out *recJ* and *xonA* while overexpressing Beta recombinase in the DH5α strain can also lead to editing efficiencies approaching 100% (14).

Researchers are also attempting to modify ncRNAs to improve editing efficiency. They explored the effects of the length of the homology arm of the donor DNA, the secondary structure of msDNA, and the strength of retron operon expression on editing efficiency. The results show that by optimizing the above factors and increasing the expression of msDNA, the editing efficiency can be significantly improved (6,13,15,16). However, there is a confusing part of the experimental results: when msDNA was expressed from the p15A (5-10 copies) plasmid, the editing efficiency is 29%, but when expressed from the pBR322 (20-40 copies) plasmid, the editing efficiency is 13%, while when the msDNA was expressed from the pUC (600 copies) plasmid (16) the editing efficiency is only 7%. This suggests that in addition to the amount of msDNA expression, there are important unknown factors that affect editing efficiency. In this study, we attempt to reveal the molecular mechanisms involved, which will provide theoretical guidance for the design and application of the retron systems.

## Results

### Construction of a retron-based gene editing system

To construct a retron-based gene editing system, we placed the retron operon containing the ncRNA coding sequence *msr/msd* and the RT coding sequence *ret* downstream of the strong constitutive promoter BBa_J23119 (https://parts.igem.org/Part:BBa_J23119). Meanwhile, the single-stranded annealing protein CspRecT was constitutively expressed to improve editing efficiency (13). The msDNA(*lacZ*_off_) expressed by the retron operon contains 92 base pairs homologous to the *lacZ* gene of the genome, which can convert the WT *lacZ* into a nonsense mutation (W17*E18*) *via* recombination. Edited colonies will appear white on LB plates containing IPTG and X-Gal (Figure 1A).

**Figure 1.**
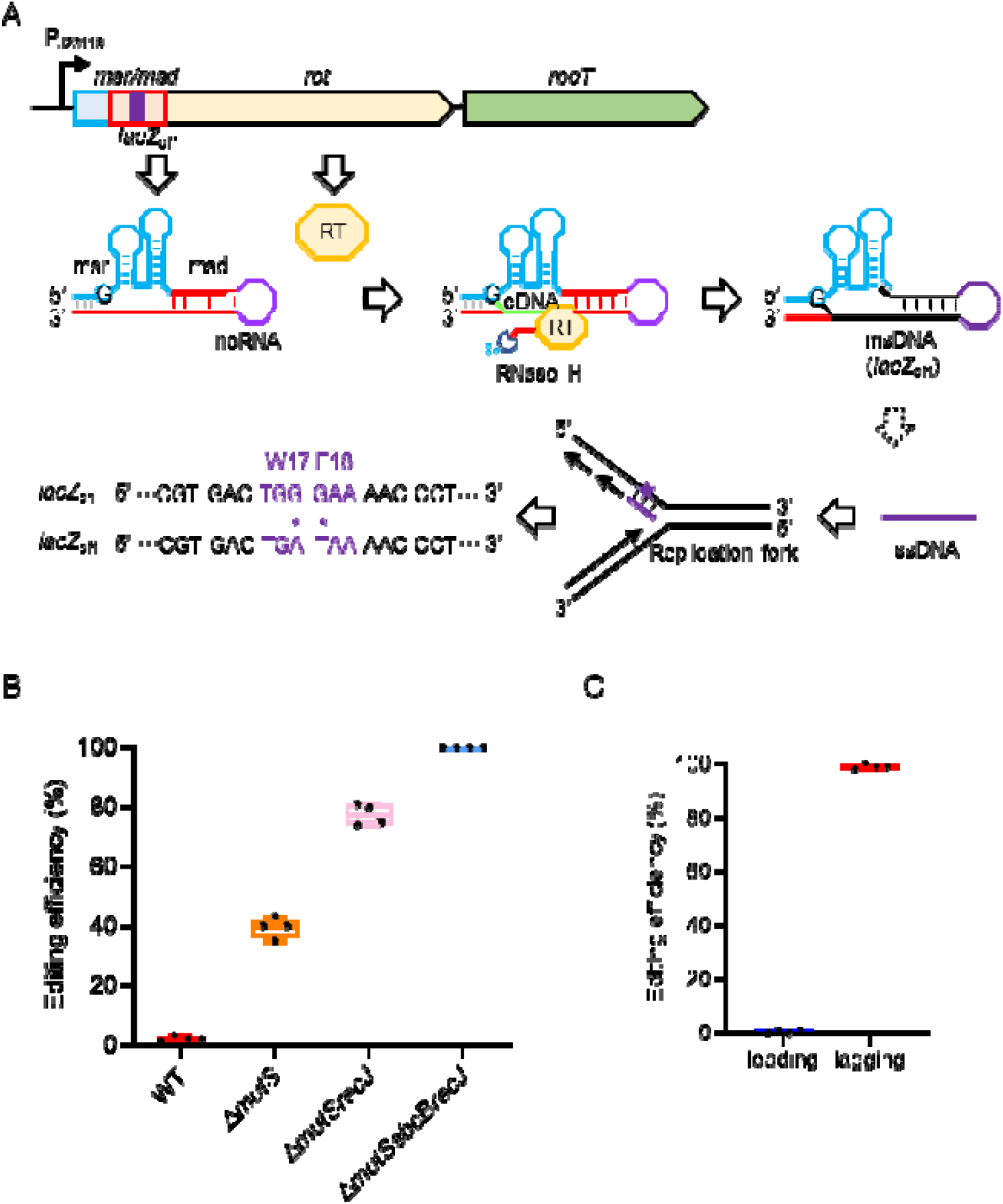
Schematic representation of the Ec86 retron system used to generate msDNA for gene editing. A. The retron operon is transcribed by the strong constitutive promoter J23119. The purple rectangle represents the inserted homologous *lacZ* sequence, which contains two mutation sites. *lacZ*_on_ represents the original *lacZ* sequence on the genome and *lacZ*_off_ represents the mutant sequence that introduces two stop codons and results in the encoding of a truncated LacZ. B. Editing efficiency of the *lacZ* gene in different *E. coli* strains. C. Editing efficiency of the *lacZ* gene when the msDNA sequence is identical to the leading and lagging strand sequences of the genome, respectively.

*E. coli* strain MG1655 and its derivatives Δ*mutS*, Δ*mutSrecJ* and Δ*mutSrecJsbcB* were transformed with retron-containing plasmids. After 24 hours of transformation, the editing efficiency was assessed by measuring the percentage of white colonies to the total number of colonies. The conversion of genomic WT *lacZ* (*lac*Z_on_) to *lacZ*_off_ in the transformants was further confirmed by colony PCR and subsequent Sanger sequencing. As shown in Figure 1B, knockdown of *mutS* increased the editing efficiency from less than 1% to 39% in MG1655, while further knockdown of *recJ* increased the editing efficiency to 78%, reaching nearly 100% in strain Δ*mutSrecJsbcB* (Figure 1B). Previous studies have suggested that msDNA with the same sequence as the Okazaki fragment (lagging strand) is more competent for gene editing (10,11). Therefore, as a comparison, we constructed another msDNA with a 92 bp *lacZ* homologous sequence (containing the *lacZ* nonsense mutation (W17*E18*)), but the msDNA sequence is identical to the leading strand in genome replication. As expected, after 24 h of incubation, the editing efficiency of msDNA-leading was less than 1%, while that of msDNA-lagging was 100% (Figure 1C). The above results confirm that the retron editing system has been successfully established.

### Effect of plasmid copy number on editing efficiency

Increasing plasmid copy number may provide more template for msDNA synthesis, thereby increasing the amount of msDNA for homologous recombination. We inserted the retron operon into plasmids with different replicons, including pSC101 (low copy), p15A (medium copy), and pUC (high copy), to explore changes in editing efficiency. The three plasmids constructed were named pSClacZ, p15AlacZ, and pUClacZ, respectively (Figure 2A). The Δ*mutSrecJsbcB* strain was transformed with above plasmids, and the editing efficiencies were recorded every 2 hours from 2 to 24 hours. As shown in Figure 2B, retrons located in different plasmids exhibited different editing efficiencies. At all time points, the editing efficiencies of Δ*mutSrecJsbcB*-p15AlacZ was the highest, approaching 100% after 12 hours of incubation, while the editing efficiency of Δ*mutSrecJsbcB*-pSClacZ continued to increase with the extension of incubation time, and the editing efficiency was higher than 90% after 24 hours of incubation. Surprisingly, the retron located on the high-copy plasmid (Δ*mutSrecJsbcB*-pUClacZ) had the lowest editing efficiency, with the highest efficiency not exceeding 20% (Figure 2B).

**Figure 2.**
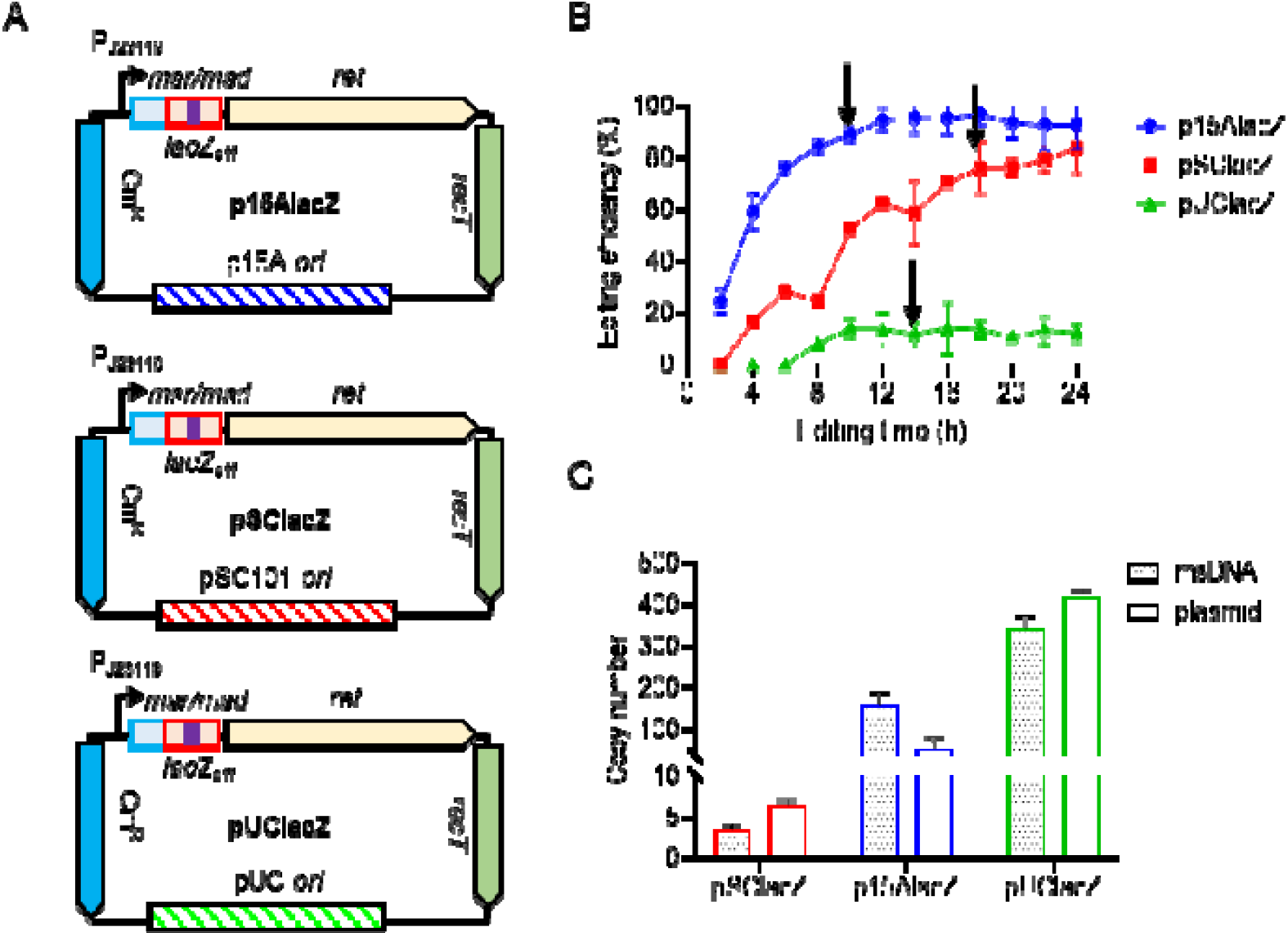
Editing efficiency of the *lacZ* gene by the retron system after altering the plasmid replicon. A. Schematic diagram of plasmids with different replicons. B. Editing efficiency of Δ*mutSrecJsbcB*-p15AlacZ, Δ*mutSrecJsbcB*-pSClacZ vs Δ*mutSrecJsbcB*-pUClacZ strains at different time points. C. Plasmid and msDNA copy numbers in Δ*mutSrecJsbcB*-p15AlacZ, Δ*mutSrecJsbcB*-pSClacZ, and Δ*mutSrecJsbcB*-pUClacZ strains measured at 10, 18, and 14 hours, respectively (indicated by arrows in B).

We then selected the time points when editing efficiency stabilized to quantify the relative abundance of each variant plasmid and msDNA. The results showed that the plasmid copy numbers of pSClacZ, p15AlacZ, and pUClacZ were approximately 6, 50, and 420, respectively, while the copy numbers of msDNA in the corresponding strains were 3, 155, and 340, respectively (Figure 2C). The above results showed that the expression amount of msDNA increased with the increase of plasmid copy number, but the increase rates of msDNA and plasmid copy number were different. More importantly, these results indicate that increased msDNA expression does not necessarily lead to improved editing efficiency.

### Plasmid *msd* sequence edited by msDNA

To elucidate the reason for the reduced editing efficiency of the Δ*mutSrecJsbcB*-pUClacZ strain, we compared the current retron editing system with an oligo-mediated editing system. We note that in the retron editing system, msDNA is synthesized from its template DNA (the *msd* region located in the retron operon), which implies that the homologous fragment msDNA has two targets in the cell: 1) Template *msd* located on plasmid; and 2) Genomic DNA. Therefore, during the editing process, the plasmid template *msd* may compete with the genomic DNA to bind the homologous fragment msDNA. If such competition exists, then an increase in plasmid copy number would result in an increase in *msd* that competes with the chromosomal target DNA, resulting in a decrease in the amount of msDNA allocated to the chromosomal target DNA, leading to a decrease in editing efficiency.

To demonstrate that msDNA can be recombined into plasmid *msd*, we developed a two-plasmid system in which one plasmid was used to generate msDNA and the other plasmid was used to observe editing. As shown in Figure 3A, the 92 bp sequence on the *kanR* gene was inserted into the *msd* region, and the *msd* sequences of pBR-P*kanX* and p15AP*kanY* contain 90 bp of homology, differing in only two bases. The Δ*mutSrecJsbcB* strain was co-transformed with pBR-P*kanX* and p15A-P*kanY* (strain 1 in Figure 3B), incubated for 24 hours, then diluted and spread on LB plates. Single colonies were randomly picked and the *msd* region on the plasmids was sequenced. The sequencing results showed a double peak at the 2 bp mutation site, indicating that the *msd* on this plasmid had been successfully edited by msDNA produced by another plasmid (Figure 3B).

**Figure 3.**
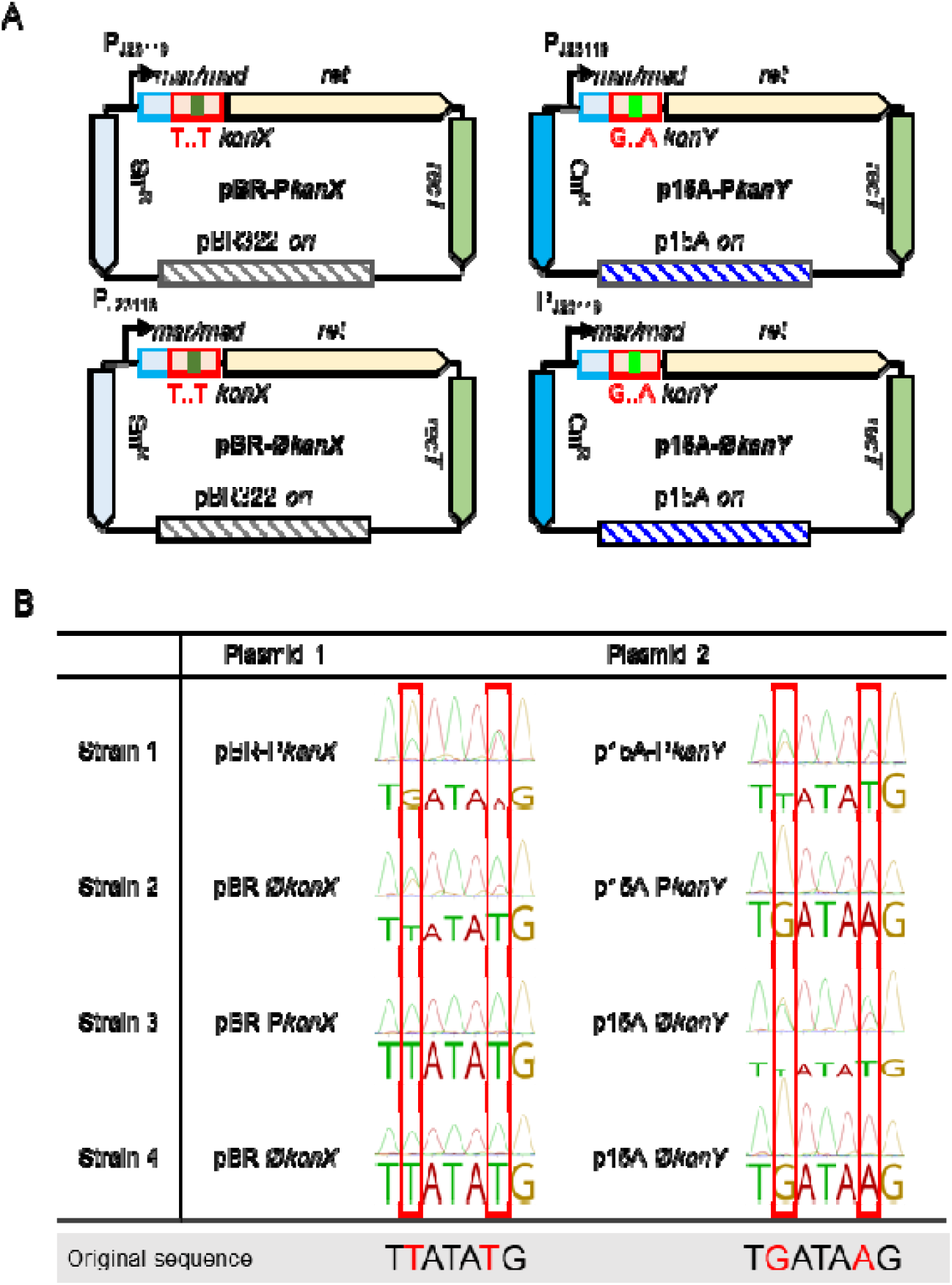
Schematic diagram of the dual plasmid system. A. Schematic diagram of four plasmids: pBR-P*kanX* and p15A-p*KanY* can generate msDNA with different *kan* sequences, respectively, while pBR-Ø*kanX* and p15A-Ø*KanY* cannot produce msDNA. B. Sequencing results of the *msd* sequences in the target region.

Next, we deleted the retron promoter and constructed pBR-Ø*kanX* and p15A-Ø*kanY* plasmids (Figure 3A). We hypothesize that the *msds* will not be edited when the retron operon is inactive. Strains 2, 3 and 4 were generated and the *msd* regions in these strains were sequenced. In strain 2, the sequencing results of the *msd* region of the pBR-Ø*kanX* plasmid showed a double peak, whereas the sequencing results of the *msd* region of the p15A-P*kanY* plasmid showed a single peak with unaltered bases (Figure 3B). Sequencing of the *msd* regions of both plasmids in strain 3 showed that pBR-P*kanX* had a single peak of unchanged bases and p15A-Ø*kanY* had a double peak (Figure 3B). In strain 4, these regions showed a single peak with unchanged bases (Figure 3B). The above results demonstrate that the msDNA synthesized can edit the *msd* on the plasmid.

### Plasmid *msd* reduced genomic DNA editing efficiency

To prove that homologous sequences on the plasmid would reduce editing efficiency, we inserted another copy of the *lacZ* homologous sequence *lacZ*_on_ into plasmid p15AlacZ and named the plasmid p15AlacZ-*lacZ*_on_ (Figure 4A). This sequence is identical to the WT *lacZ* sequence on the genome but differs by 2 bp from the *lacZ*_off_ sequence in the retron operon.

**Figure 4.**
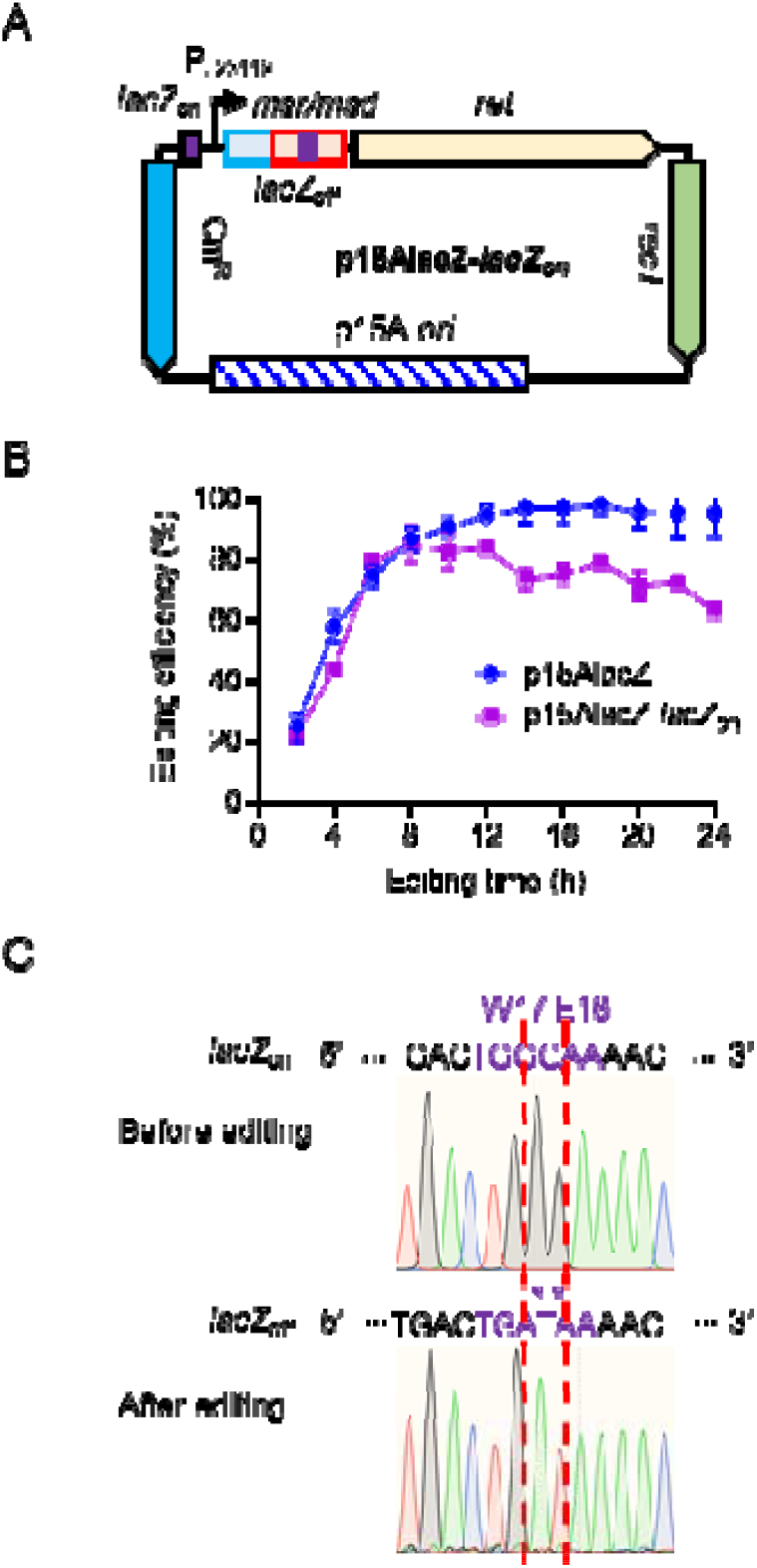
Additional copies of *msd* reduce *lacZ* gene editing efficiency. A. Schematic diagram of plasmid p15AlacZ-*lacZ*_on_. The inserted homologous *lacZ* sequence *lacZ*_on_ is indicated as a purple rectangle. B. Editing efficiency of Δ*mutSrecJsbcB*-p15AlacZ, Δ*mutSrecJsbcB*-p15AlacZ-*lacZ*_on_ strains at different time points. C. Sequencing results of the homologous *lacZ* sequences on the plasmids before and after editing.

Continuous monitoring of editing efficiency showed that from hour 12 onwards, the editing efficiency of Δ*mutSrecJsbcB*-p15AlacZ-*lacZ*_on_ was significantly lower than that of strain Δ*mutSrecJsbcB*-p15AlacZ (Figure 4B). A 24-hour culture of the Δ*mutSrecJsbcB*-p15AlacZ-*lacZ*_on_ strain was diluted and plated, and colonies were randomly selected and sequenced. The sequencing results showed that the sequence GG of the *lacZ*_on_ region on the plasmid was mutated to AT (Figure 4C), proving that *lacZ*_on_ was edited by msDNA. Based on these results, we conclude that the presence of homologous sequence on plasmid compete with homologous sequence on genomic DNA, thereby reducing retron editing efficiency.

### msDNA served as the leading strand during plasmid replication effectively improved genomic DNA editing efficiency

The pUC plasmid has a ColE1-based origin (*ori*) and is a θ-type replication plasmid (17). Previous studies have also shown that plasmids with a ColE1 origin have the potential to replicate to either side of the *ori* at different rates, but the precise replication termination site (*ter*) is unknown (18,19). Given that the lagging strand msDNA is more competent of editing than the leading strand msDNA (9-11), we hypothesized that when msDNA serves as the leading strand of the plasmid, the editing effect of msDNA on *msd* will be diminished (Figure 5A). In this way, *msd* will not be able to compete with homologous genomic DNA sequences, and the editing efficiency of genomic DNA will be improved.

**Figure 5.**
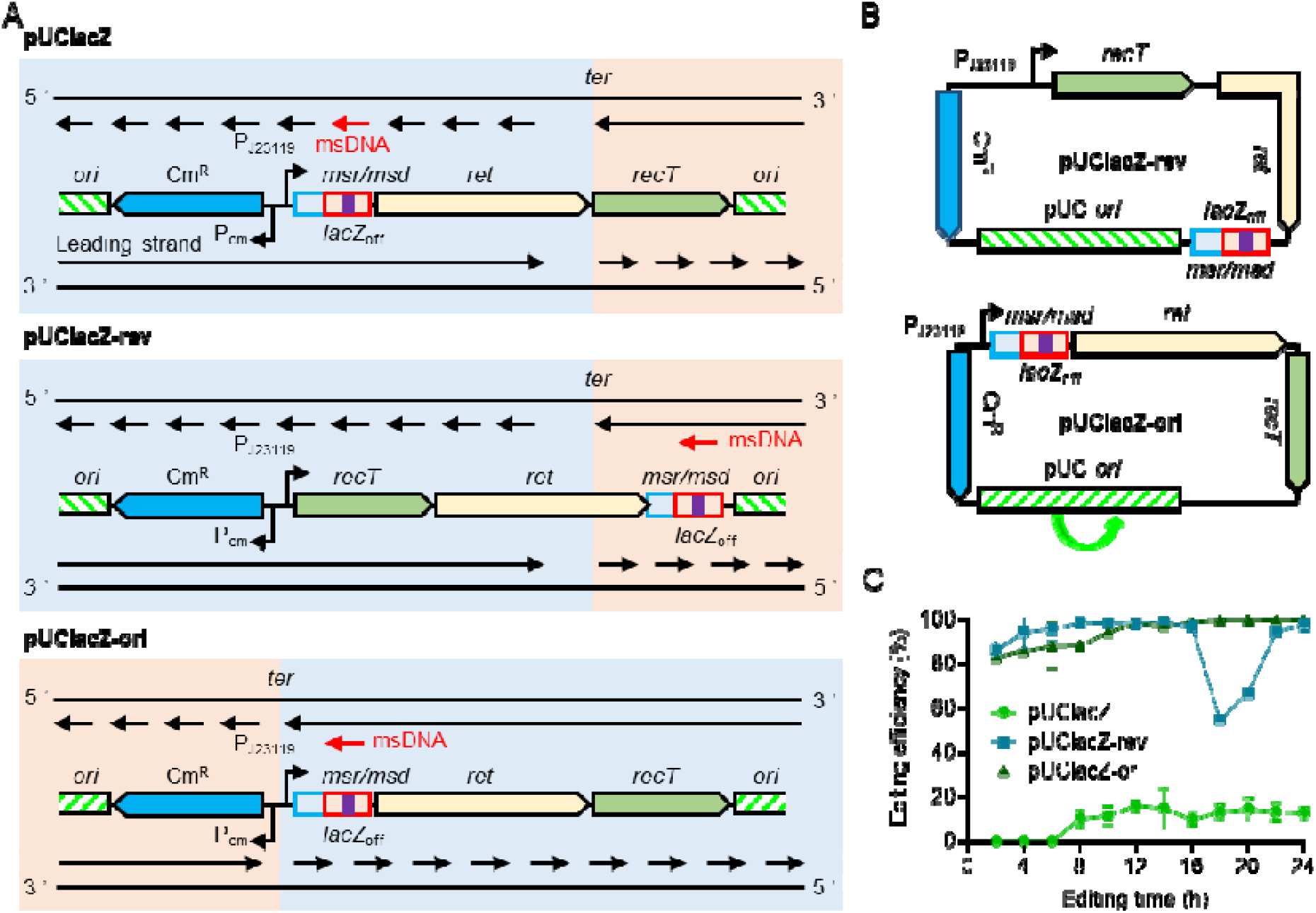
Editing efficiency of *lacZ* gene after relocating the *msd* position relative to the *ori*. A. Schematic diagram of the theta (θ) type replication in plasmids pUClacZ, pUClacZ-rev and pUClacZ-ori: DNA unwinds at the *ori* site from where the replication begins. Two replication forks start at *ori* and then travel in opposite directions until they meet at *ter*. Black lines above and below the operon: parental (template) strands; short black arrows: lagging strand; long black arrows: leading strand. short red arrows: homologous sequences present on msDNA; msDNA efficiently edits the *msd* (pUClacZ), msDNA is unable to edit the *msd* (pUClacZ-rev and pUClacZ-ori). Blue and pink shaded area: replication fork extension region, DNA replication is faster in the blue region than in the pink region. B. Schematic diagram of the pUClacZ-rev and pUClacZ-ori plasmids. C. Editing efficiency of Δ*mutSrecJsbcB*-pUClacZ, Δ*mutSrecJsbcB*-pUClacZ-rev and Δ*mutSrecJsbcB*-pUClacZ-ori strains at different time points.

Based on the above speculation, we first reversed the positions of *msr*/*msd, ret* and *recT* on the pUClacZ plasmid and constructed a new plasmid, pUClacZ-rev (Figure 5B). On the pUClacZ-rev plasmid, the *ori* and transcription direction of the retron operon remain unchanged, but the order of the gene changes. In this way, we move the location of *msd* from an area that can be edited by msDNA to an area that cannot be edited (Figure 5A). The Δ*mutSrecJsbcB* strain was transformed with pUClacZ-rev and pUClacZ, respectively, and the editing efficiencies were recorded every 2 hours until incubation for 24 hours. The results showed that after only 2 hours of incubation, the editing efficiency of Δ*mutSrecJsbcB-*pUClacZ-rev exceeded 80%, at 6 hours, editing efficiency was close to 100%, which was significantly higher than that of Δ*mutSrecJsbcB-*pUClacZ (Figure 5C). We also observed an interesting phenomenon that after 18 hours of incubation, the editing efficiency of Δ*mutSrecJsbcB-*pUClacZ-rev dropped significantly to only 55%, and then gradually recovered to nearly 100% by 22 hours. The mechanisms underlying this phenomenon need to be further explored.

In another attempt to improve editing efficiency, we reversed the orientation of the *ori* on the pUClacZ plasmid to generate the plasmid pUClacZ-ori. We hypothesized that the *msd* would change from an editable region to a non-editable region due to the different replication rates of the plasmid from both sides (Figure 5A). The editing efficiency recorded for the Δ*mutSrecJsbcB*-pUClacZ-ori strain showed that it indeed edited the genome *lacZ* with considerable efficiency, exceeding 80% at 2 hours and close to 100% at 10 hours (Figure 5C). Through the above experiments, we demonstrated that reducing msDNA editing of *msd* is indeed an effective means to improve the efficiency of genomic DNA editing.

In addition, we tried to insert a 4.2 kb long dCas9 coding sequence on the left side of the *ori* of pUClacZ which is only 4.0 kb in length. We expected dCas9 coding sequence to push the *msd* to non-editable region. The Δ*mutSrecJsbcB* strain was transformed with the 8.2 kb-long plasmid pUClacZ-long, and the editing efficiency was recorded. Unexpectedly, the Δ*mutSrecJsbcB*-pUClacZ-long strain was almost unable to edit genome *lacZ*, with an editing efficiency of less than 1% (data not shown). Plasmid sequencing showed that this was due to a large number of deletions on the plasmid, resulting in an ineffecitve retron. From this result we can learn that plasmid stability is also a very important consideration when using the retron system for gene editing.

## Discussion

In this study, we deeply studied the effect of msDNA template *msd* on editing efficiency and found that the increase of *msd* increased msDNA expression and competed more for msDNA bound to genomic DNA. Previous studies have focused more on the synthesis of msDNA and its improved efficiency in participating in the DNA replication process, but less on the impact of the amount of *msd* on editing efficiency. Both gene editing based on homologous recombination and gene editing based on CRISPR-Cas combined with λ-Red recombination require the introduction of homologous fragments of the target gene into the strain (20,21). There are two main methods for introducing homologous fragments: one is to introduce oligonucleotides by electroporation (22);the other is to clone the homologous fragments into plasmids and introduce them as plasmid fragments. In the above editing methods, there are two elements, namely the homologous fragment, and the object to be edited (genomic DNA). However, in the retron editing system, homologous fragments are synthesized spontaneously within the cell using *msd* as a template (6,13,14,23). This means that the retron editing system contains three elements: homologous fragment template (*msd*), homologous fragment (msDNA), and the object to be edited (genomic DNA). Therefore, two targets of homologous fragments exist in the cell: 1) the homologous fragment template *msd* located on the plasmid, and 2) the genomic DNA. This implies that during the editing process, the plasmid template *msd* may compete with genomic DNA for binding to the homologous fragment msDNA. Because different kinds of plasmids have different copy numbers, there are differences in the ability of different plasmids to compete with genomic DNA. We believe that this process cannot be ignored; first, it may have an important impact on the efficiency of gene editing; second, in retron-based directed evolution systems, for example, msDNA edits *msd* after mutations are introduced into msDNA via error prone T7 RNA polymerase, thereby producing a heritable mutation.

## Materials and Methods Plasmid construction

Plasmids were constructed using pFF745 as the parent plasmid (5). Primer pairs used for plasmid construction were synthesized by Tianyi Huiyuan (Wuhan, Hubei, China). All plasmids generated in this study were assembled using the Hieff Clone Plus multi one-step cloning kit (Yeasen, Shanghai, China) and confirmed by sequencing (Quintarabio, Wuhan, Hubei, China). The constructed plasmids were transformed into bacterial cells using the calcium chloride (CaCl_2_) method to generate corresponding derivative strains.

### Bacterial strains and growth conditions

*E. coli* DH5a was used for all cloning experiments. Δ*mutS*, Δ*mutSrecJ* and Δ*mutSrecJsbcB* strains were generated via the CRISPR-Cas9 System (24). *E. coli* strain MG1655 containing the corresponding plasmids (as shown in the figures) and its derivatives Δ*mutS*, Δ*mutSrecJ* and Δ*mutSrecJsbcB* were grown at 37°C in LB medium containing 10 g tryptone, 5 g yeast extract, and 10 g NaCl per L of water and supplemented with chloramphenicol (15 μg/mL) or Spectinomycin (20 μg/mL) if necessary.

### Editing efficiency quantification

We calculated gene editing efficiency by blue-white screening, i.e., we could know the editing efficiency by counting the proportion of white colonies to the total number of colonies. The corresponding plasmid (about 200 ng) was electroporated into 200 μL Δ*mutSrecJsbcB* competent cell, and incubated with 800 μL LB medium at 37 °C for 1 h. After the incubation, we inoculated 1 mL of culture solution into 50 mL of LB medium containing chloramphenicol, incubated it at 37 °C at 200 r/min until a specific time, diluted the bacterial solution to an appropriate concentration, and plated it on LB containing IPTG, X-gal and chloramphenicol. The plates were incubated at 37 °C for 24 hours and the number of white and blue colonies was counted.

### Determination of plasmid and msDNA copy numbers

We placed 100 μL of logarithmic growth phase cells in a boiling water bath and heated them for 10 minutes for cell lysis. Lysates containing genomic DNA, plasmid DNA, and msDNA were stored at −20°C until use, and qRT-PCR was used to quantify the copy number of plasmid and msDNA (6) using three sets of primers with the sequences of tdk-qPCR-F (5’-CCGCAATGAATGCGGGGTAAG-3’) (5’-GCAGGCGATGACAAACCTATACG-3’), and ret-qPCR-F tdk-qPCR-R (5’-AGGCGACTCGCATATCTGTTG-3’) and ret-qPCR-R (5’-CGTAGAACCCATCCTTGTAAGGC-3’), msd-qPCR-F (5’-CTGAGTTACTGTCTGTTTTCCTG-3’) and msd-qPCR-R (5’-TCAGAAAAAACGGGTTTCCTGAATTC-3’). The first two sets of primers were used to amplify genomic DNA and plasmid DNA, respectively, and the obtained cycle thresholds (Ct) were named C_tg_ and C_tp_, respectively. The plasmid copy number was calculated according to the following formula: plasmid copy number = 2^-ΔCt^ = 2 ^(Ctg - Ctp)^. The third set of primers was designed in the msr/msd region to amplify not only msDNA but also msr/msd on the plasmid. Therefore, the obtained cycling threshold C_tm_ is the result of the co-amplification of the plasmid and msDNA template. Furthermore, msDNA copy number can be calculated by the following formula: msDNA copy number = 2 ^(Ctg - Ctm)^ - plasmid copy number.

qRT-PCR was carried out on a Bio-Rad CFX96 real-time quantitative PCR instrument, with a reaction volume of 20 uL, including 10 uL of Hieff® qPCR SYBR Green Master Mix (Yeasen, China), and 1 uL each of upstream and downstream primers (final concentration is 0.5 umol/L), 0.4 uL of template DNA, and H_2_O to a final volume of 20 uL. The following conditions were used for the qRT-PCR reaction: (1) pre-denaturation at 95 °C for 4 min for cyclic reactions (40×) (2) denaturation at 95 °C for 10 s and (3) amplification at 60 °C for 30s. Fluorescence readings were taken and data were recorded during the amplification step, and a melting curve was plotted at the end of 40 cycles. Amplification curves and melting curves were plotted by CFX Maestro software 2.3 (Bio-Rad Laboratories, TX, USA). For data analysis, technical and biological triplet data were obtained. The expression of each sample is presented as mean±SD. The values were subjected to t-test, and GraphPad Prism 9.0 was used for t-tests and graph drawing.

## Acknowledgments

This work was supported by the National Natural Science Foundation of China (31971339 and 32171422), the National Key Research and Development Program of China (2022YFF1000700), and the Fundamental Research Funds for the Central Universities (2662022SKYJ004). This work was also supported by the Science and Technology Research Project of Jiangxi Provincial Department of Education (Grant No: GJJ211722).

